# Cell-matrix adhesion controls Golgi organization and function by regulating Arf1 activation in anchorage dependent cells

**DOI:** 10.1101/261842

**Authors:** Vibha Singh, Chaitanya Erady, Nagaraj Balasubramanian

**Keywords:** Golgi apparatus, matrix, adhesion, Arf1, glycusylation

## Abstract

Cell-matrix adhesion regulates membrane trafficking to control anchorage-dependent signaling. While a dynamic Golgi complex can contribute to this pathway, its control by adhesion remains untested. We find the loss of adhesion rapidly disorganizes the Golgi in mouse and human fibroblast cells, its integrity restored rapidly on re-adhesion to fibronectin (but not poly-l-lysine coated beads) along the microtubule network. Adhesion regulates the trans-Golgi more prominently than the cis /cis-medial Golgi, though they show no fallback into the ER making this reorganization distinct from known Golgi fragmentation. This is controlled by an adhesion-dependent drop and recovery of Arf1 activation, mediated through the Arf1 GEF BIG1/2 over GBF1. Constitutively active Arf1 disrupts this regulation and prevents Golgi disorganization in non-adherent cells. Adhesion regulates active Arf1 binding to the microtubule minus-end motor protein dynein to control Golgi reorganization, which ciliobrevin blocks. This regulation by adhesion controls Golgi function, promoting cell surface glycosylation on the loss of adhesion that constitutively active Arf1 blocks. This study hence identifies cell-matrix adhesion to be a novel regulator of Arf1 activation, controlling Golgi organization and function in anchorage-dependent cells.

**Summary Statement:** This study identifies a role for cell-matrix adhesion in regulating organelle (Golgi) architecture and function which could have implications for multiple cellular pathways and function.

## INTRODUCTION

Cell-matrix adhesion is a vital regulator of many cellular processes (Berrier and Yamada, 2007; Caswell et al., 2009; Danen and Yamada, 2001) and disease conditions (Desgrosellier and Cheresh, 2010; Wu and Reddy, 2012). A role for integrin-mediated adhesion in regulating membrane trafficking is seen to affect membrane order (Gaus et al., 2006), receptor mobility and activation to drive anchorage-dependent signalling(Balasubramanian et al., 2007a, 2010; Pawar et al., 2016). Loss of adhesion dramatically turns off this signalling that recovers on re-adhesion to matrix proteins(del Pozo et al., 2004a; Schwartz, 1997). Cancer cells deregulate this trafficking to become anchorage-independent, supporting oncogenic transformation (Guo and Giancotti, 2004; Schwartz, 1997). An important mediator of trafficking and processing of membrane lipids and proteins in the cell is the Golgi apparatus (Rodriguez-Boulan and Müsch, 2005). Known to be a dynamic structure, the organization, and positioning of the Golgi in the cell is vital to directional trafficking and secretion during processes like cell polarization, migration, and division (Wilson et al., 2011; Yadav and Linstedt, 2011). Integrin-mediated cell adhesion is seen to regulate these cellular process, though its role in controlling Golgi architecture and function remains largely untested.

The architecture of the Golgi apparatus is subject to many variables and directly contributes to Golgi inheritance and function (Shorter and Warren, 2002). In mammalian cells, the Golgi is made up of a series of flattened cisternal stacks that are not homogeneous and contain different resident proteins and enzymes that allow the Golgi to be structurally divided into cis, medial, and trans regions (Shorter and Warren, 2002). The cis Golgi receives newly synthesized proteins and lipids from the Endoplasmic Reticulum (ER), while its cargo exits from the trans-Golgi (Glick and Luini, 2011). The Trans Golgi network (TGN) is the main cargo sorting compartment where proteins and lipids are sorted and delivered to their destinations. The Golgi typically has a perinuclear localization around the microtubule organizing center (MTOC) (Hurtado et al., 2011), its structure being controlled by both the microtubule(Sandoval et al., 1984) and actin cytoskeleton (Valderrama et al., 1998). It undergoes dramatic fragmentation during cell division (Colanzi and Corda, 2007), apoptosis (Hicks and Machamer, 2005), and disease conditions like neuronal degeneration (Nakagomi et al., 2008) and cancers (Petrosyan, 2015). This fragmentation is at times irreversible, like during apoptosis and sometimes reversible, as seen during cell division.

The Golgi complex is held together and reorganized along the microtubule network(Sandoval et al., 1984) with the help of membrane stacking proteins (GRASP65, GRASP55) differentially localised to the distinct Golgi compartments (Vinke et al., 2011). Small GTPase Arf1, acts as a major regulator of Golgi structure and function. Upon its activation Arf1 associates with Golgi membranes and is released from Golgi membranes into the cytosol on its inactivation (Donaldson et al., 2005). This association and dissociation of Arf1 is regulated by Golgi-associated, guanine nucleotide exchange factors (GEFs) and GTPase activating proteins (GAPs). Arf1 binds and/or regulates adaptor, stacking, structural proteins and lipid-modifying enzymes, phospholipase D (PLD) and phosphatidylinositol 4-phosphate 5-kinase (PIP5-kinase) on the Golgi membrane (Donaldson et al., 2005). It is hence able to influence multiple aspects of Golgi structure and function in cells.

Integrin-mediated adhesion is seen to regulate Arf family GTPase Arf6, through the Arf6 GEF ARNO (Lee et al., 2014; Pawar et al., 2016), to regulate adhesion-dependent membrane exocytosis and signaling(Balasubramanian et al., 2007a, 2010; Pawar et al., 2016). Adhesion is also seen to control cell cycle progression (Ben-Ze’ev and Raz, 1981; Schwartz and Assoian, 2001) and apoptosis (Meredith and Schwartz, 1997), these regulatory pathways are bypassed by cancer cells to drive anchorage-independent signaling and growth (Guo and Giancotti, 2004; Schwartz, 1997). With the differential organization of the Golgi is seen to be associated with normal and cancer cell function, the possible role integrin-mediated adhesion could have in regulating the same remains unexplored. In this study, we hence tested and identify the presence of such a regulatory pathway that drives cell adhesion-dependent Golgi organization and function in anchorage-dependent cells.

## RESULTS

### Cell adhesion regulates Golgi organization

Stable adherent (SA) wild type mouse embryonic fibroblasts (WT-MEFs) with a classical organized Golgi phenotype, grown in low serum conditions for 12 hours to suppress growth factor signalling, when detached (5 min for processing) and held in suspension with 1% methylcellulose for 120 mins rapidly disorganize their cis-Golgi (detected with anti GM130 antibody staining) (Fig. 1a), cis-medial Golgi (detected with Mannosidase II-GFP) (Supplementary Fig. 1b) and most prominently the trans-Golgi (detected with GalTase-RFP) (Fig. 1b). Re-adhesion of cells on fibronectin-coated coverslips for 5 mins while not significantly affecting cell shape or volume (Supplementary Fig. 1a) dramatically restores cis (Fig. 1a), cis-medial (Supplementary Fig. 1b) and trans-Golgi organization (Fig. 1b). Percentage distribution of the organized vs disorganized trans-Golgi phenotype when compared in the cell population confirms the disorganized phenotype to be prominent in suspended cells, reversed rapidly on re-adhesion (Supplementary Fig. 1e). This reorganization of the Golgi is further reflected in a significant reduction in the number of cis and trans-Golgi objects in re-adherent cells relative to when they are detached (5’) and suspended (120’) (Fig. 1c, 1d, Graph in Supplementary Fig. 1b). The net cis-Golgi volume, when compared across suspended and re-adherent cells, did not change significantly (Supplementary Fig. 1f). The trans-Golgi that undergoes a more extensive reorganization did show an increase in net Golgi volume on the loss of adhesion, significantly reduced on re-adhesion (Supplementary Fig. 1g). We further looked at the co-localization of the cis, cis-medial and trans-Golgi relative to each other and find the loss of adhesion preferentially disorganizes the trans-Golgi. This is reflected in the number of trans Golgi objects being significantly more than cis or cis-medial Golgi (Fig1c, 1d, Graph in Supplementary Fig 1b) and their colocalization (Pearson coefficient) being significantly low in suspended cells, restored as the Golgi reorganizes on re-adhesion (Fig 1f, top and middle panel). In contrast, the cis and cis medial Golgi show significant overlap even when they are disorganized in suspended cells, their colocalization only marginally better in re-adherent cells (Fig 1f, lower panel). Interestingly, on the loss of adhesion, the disorganized cis Golgi (GM130) failed to show any significant overlap with the ER (KDEL-RFP) (Fig 1g, 1h), suggesting this phenotype to be distinct from known Golgi fragmentation seen to cause fall back of the cis Golgi to the ER (Mardones et al., 2006).

**Figure 1.**
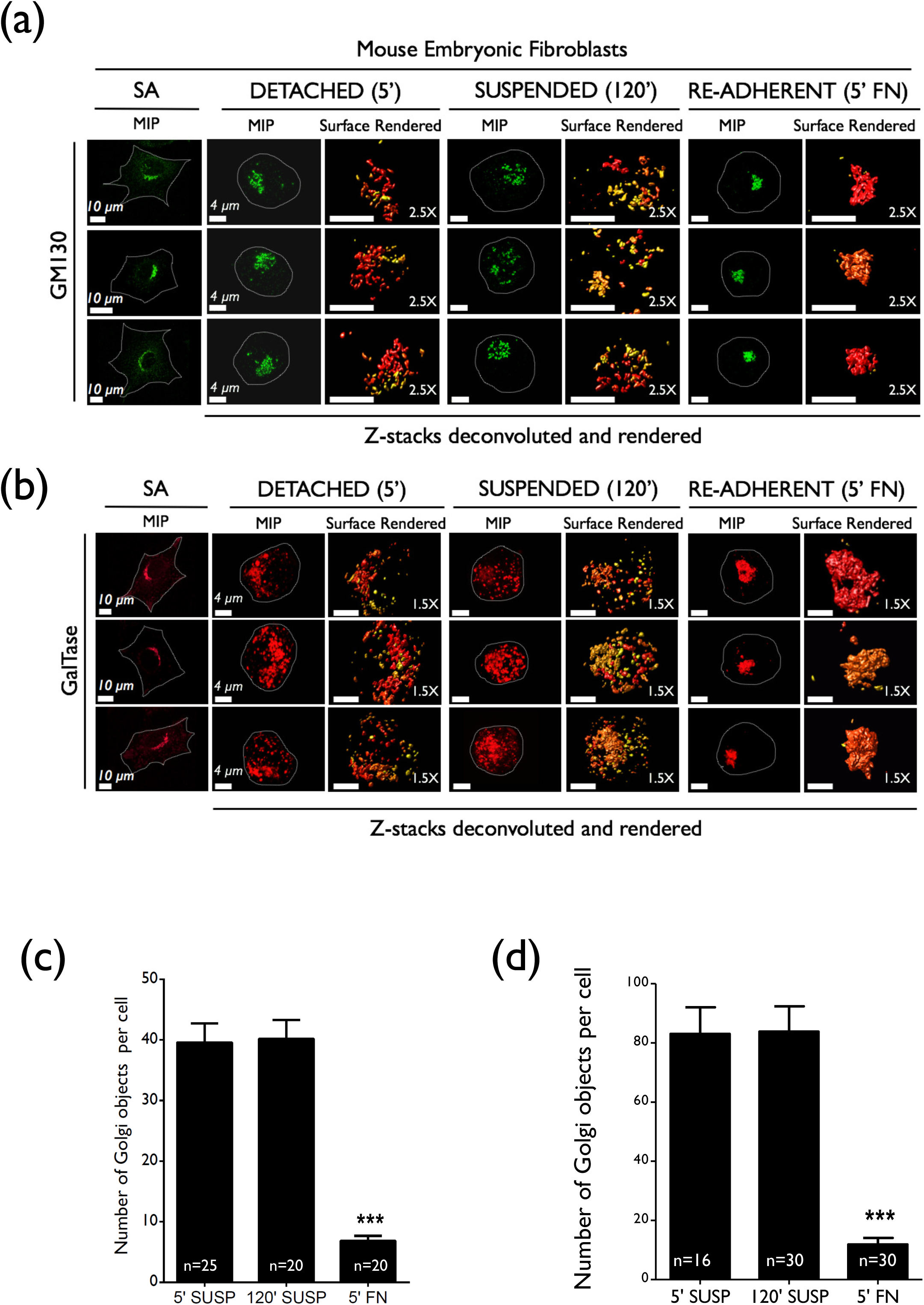
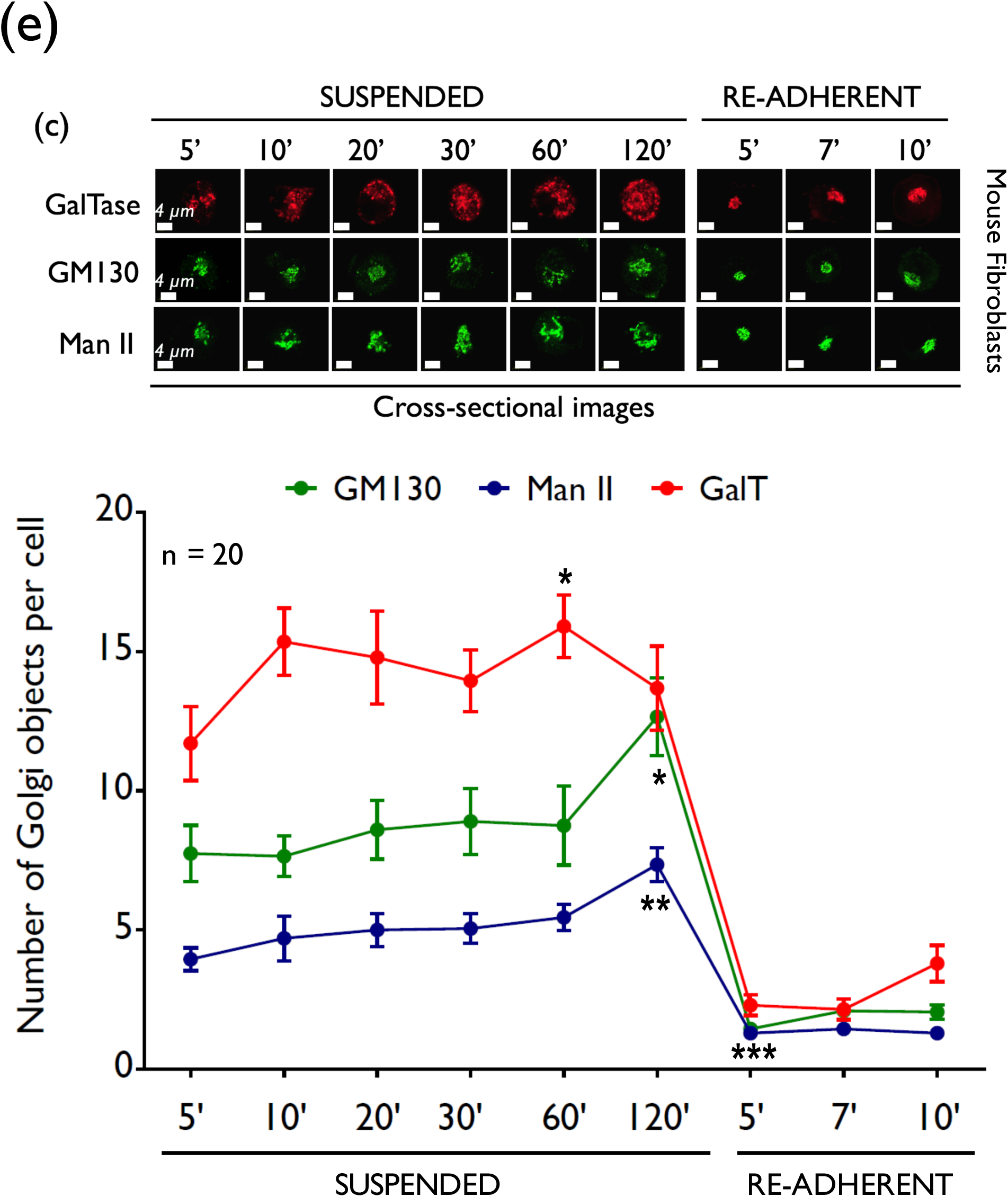
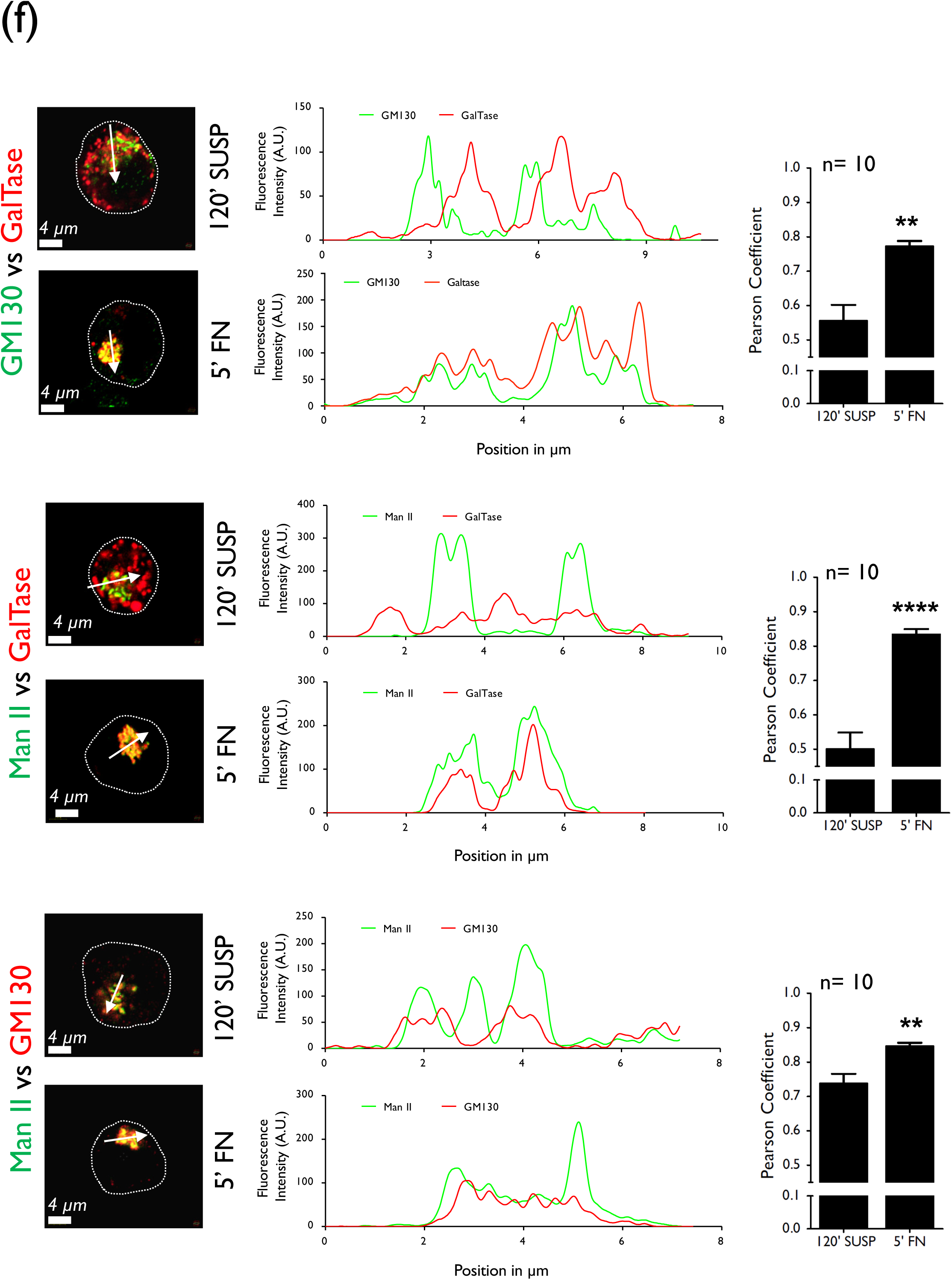
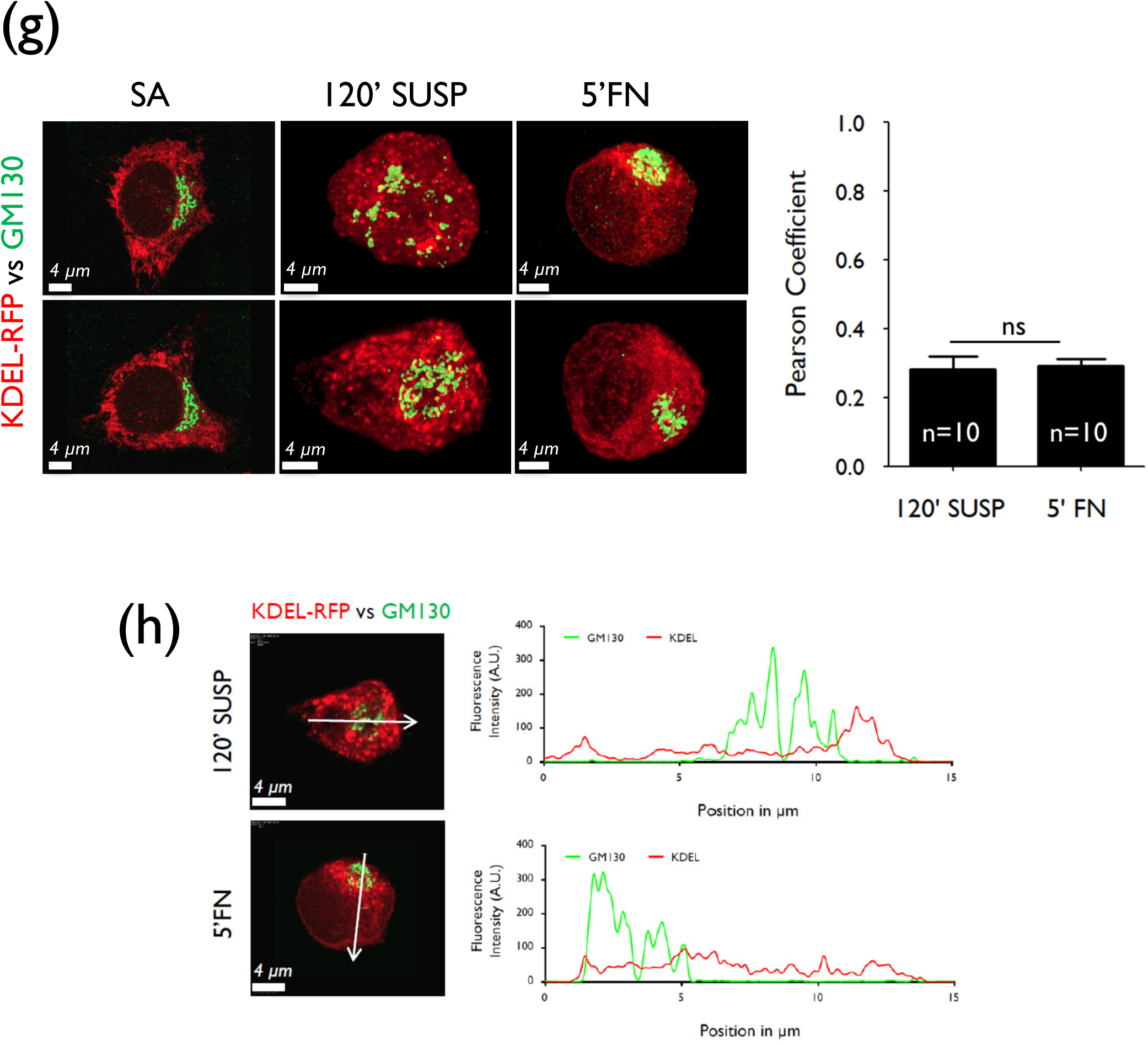
Adhesion regulates Golgi organization. WT-MEFs (**a**) un-transfected or (**b**) transfected with GalTase RFP serum starved for 12 hours as stable adherent (SA) were detached (5’) and held in suspension for 120 mins (120’) and re-plated on fibronectin for 5 mins (5’ FN). Their (**a**) cis Golgi was detected by anti-GM130 antibody staining (GM130) and (**b**) trans Golgi using the GalTase RFP localization (GalTase). Maximum intensity projections (MIP) of de-convoluted confocal images for 3 representative cells for GM130 and GalTase RFP for each time point are shown (left panel at each time point). De-convoluted confocal Z stacks were surface rendered and zoomed for clarity (2.5X for GM130 and 1.5X for GalTase) (right panel at each time point). Stable adherent cells (SA) are represented only as maximum intensity projections (MIP). Following deconvolution of detached (5’ SUSP), suspended (120’ SUSP) and re-adherent (5’ FN) cells discontinuous Golgi objects per cell detected labeled for (**c**) GM130 or (**d**) GalTase were determined using the Huygens image analysis software. The graph represents mean ± SE from a minimum of 16 cells and maximum of 30 cells (as indicated in each bar) from three independent experiments. Statistical analysis comparing 120’ SUSP and 5’ FN was done using the Mann Whitney’s test (*** p value <0.0001) (**e**) Stable adherent WT-MEFs, expressing GalTase-RFP (top panel), ManII-GFP (lowest panel) or stained with anti-GM130 antibody (middle panel) were detached and held in suspension for increasing time points 5 mins to 120 mins (5’, 10’, 20’, 30’, 60’, 120’ Suspension) and then re-plated on FN for increasing times (5’, 7’, 10’ Re-adherent). Cross-sectional confocal images for trans-Golgi (GalTase), cis Golgi (GM130) and cis medial Golgi (Man II) were deconvoluted and discontinuous Golgi objects per cell determined using the Huygens image analysis software. Representative cross-section images for each Golgi marker across suspension and re-adhesion time points are shown in the panel above graph. The graph represents mean ± SE for Golgi objects in 20 cells from two independent experiments. Statistical analysis comparing object numbers in cells suspended for 5 min to 60 mins and 120 mins was done using the Mann Whitney’s test. This analysis was extended to compare the decrease in object numbers between cells suspended for 120 mins and re-adherent on fibronectin for 5 mins (* p-value <0.05, ** p-value <0.01, *** p-value <0.0001). Scale bar in all images is set to 4 µm. (**f**) Serum-starved WT-MEFs transiently expressing or immunostained with a combination of Golgi markers, GM130 vs. GalTase (top panel), Man II vs. GalTase (middle panel), and Man II vs GM130 (bottom panel) were held in suspension for 120 mins (120’ SUSP) and re-plated on fibronectin for 5 mins (5’ FN). Fluorescence intensity measurements made for these Golgi markers along a line (marked by arrow) in representative 120’ SUSP and 5’ FN re-adherent cell were plotted and compared. Markers labeled in green or red are represented in a line of the same color in the graph. Co-localization in the entire confocal cross-sections were determined with Huygens software and Pearsons coefficient represented in the graph (mean ± SE) (10 cells from one experiment). Scale bar in all images is set at 4 µm. Statistical analysis done using the Mann Whitney’s test (** p-value <0.01, **** p-value <0.00001). (**g**) Representative images of serum-starved WT-MEFs expressing ER marker KDEL (KDEL-RFP) were stable adherent (SA), held in suspension for 120 mins (120’ SUSP), re-plated on fibronectin for 5 mins (5’ FN) and cis Golgi stained with anti GM130 antibody (GM130). Co-localization for ER (KDEL-RFP) and Golgi (GM130) in the entire confocal cross-sections were determined with Huygens software and Pearsons coefficient represented in graph (mean ± SE) (**h**) Colocalization was also assessed in a representative 120’ SUSP and 5’ FN re-adherent cell, using a line plot across the cell (marked by arrow) for each marker using Huygens image analysis software. Fluorescence intensity measurements obtained were plotted for both the markers and their profiles compared. All scale bars in images are 4 µm, except stable adherent cells set at 10µm.

We also tested the kinetics of reorganization for the cis, cis-medial and trans Golgi and find loss of adhesion in the first 5 minutes (needed for processing of detached cells) to significantly disorganize the Golgi, with the cis/cis-medial and trans Golgi object numbers increasing only marginally over time in suspension (Fig 1e). Re-adhesion of cells to fibronectin for 5 mins (minimum time needed for sufficient cells to be re-adherent for imaging) shows the Golgi to be reorganized, reflected in a significant decrease in cis, cis-medial and trans-Golgi objects (Fig 1e). The rapid nature of this reorganization on re-adhesion to fibronectin in lower serum conditions supports a role for integrin-mediated adhesion in driving this pathway. Loss of adhesion and re-adhesion under these conditions is seen to regulate Akt activation (del Pozo et al., 2004a) (Supplementary Fig. 1c) and GM1 endocytosis (Balasubramanian et al., 2007a) (Supplementary Fig. 1d) in these cells, both known to be controlled by integrins.

Such an adhesion-dependent regulation of the Golgi was also observed using cis-medial (ManII GFP) and trans (GalTase RFP) Golgi markers in anchorage-dependent human foreskin fibroblasts (Supplementary Fig. 1i) and human endothelial cells, EA.hy926 (data not shown). A similar and significant difference in trans vs cis-medial Golgi object numbers when suspended was observed, restored on re-adhesion (graphs in Supplementary Fig. 1i), confirming their differential regulation in these cells (like noted for mouse fibroblasts). This suggests adhesion-dependent regulation of the Golgi to be conserved across cell types.

### Fibronectin vs poly-L-lysine coated beads differentially restore Golgi organization

The rapid nature of adhesion mediated regulation of the Golgi organization does suggest early adhesion events to drive the same. This leads us to test if in non-adherent serum starved WT-MEFs, binding to fibronectin (FN-Bead) vs poly-L-lysine (PLL-Bead) coated polystyrene beads affects Golgi architecture. Fibronectin coated beads dramatically restore the trans-Golgi (GalTase RFP) organization, relative to control and poly-L-lysine coated beads (Fig 2a). This is reflected in a significant drop in trans Golgi object numbers in FN bead-bound cells (Graph in Fig 2a). Percentage distribution of the organized vs disorganized trans-Golgi phenotype across cells further confirms the effect fibronectin beads have over control and PLL beads (Fig. 2b). The differential effect Fibronectin vs PLL beads have suggested variable binding and clustering of integrins (Tran et al., 2002) and resulting downstream signaling drives the Golgi organization. PLL coated beads likely cause some integrin activation (relative to control) that reflects in a small reduction in Golgi object numbers (Graph in Fig 2a), but is not prominent enough to affect the distribution profile for the phenotype in cells (Graph in Fig 2b). Fibronectin coated beads are also seen to support the enrichment of GM1 labelled raft microdomains (detected with Cholera Toxin B) in cells, confirming their spatial activation of integrins (Supplementary Fig. 2a). With a known integrin-growth factor crosstalk reported in WT-MEFs, we also tested if the presence of serum growth factors (5% FBS) affects adhesion-dependent Golgi architecture. In cells with serum (5% FBS), like with cells in low serum (0.2% FBS) conditions, the Golgi similarly disorganizes on the loss of adhesion and recovers on re-adhesion. This further suggests adhesion be a primary regulator of Golgi organization in WTMEFs (Supplementary Fig. 2d, 2e).

**Figure 2.**
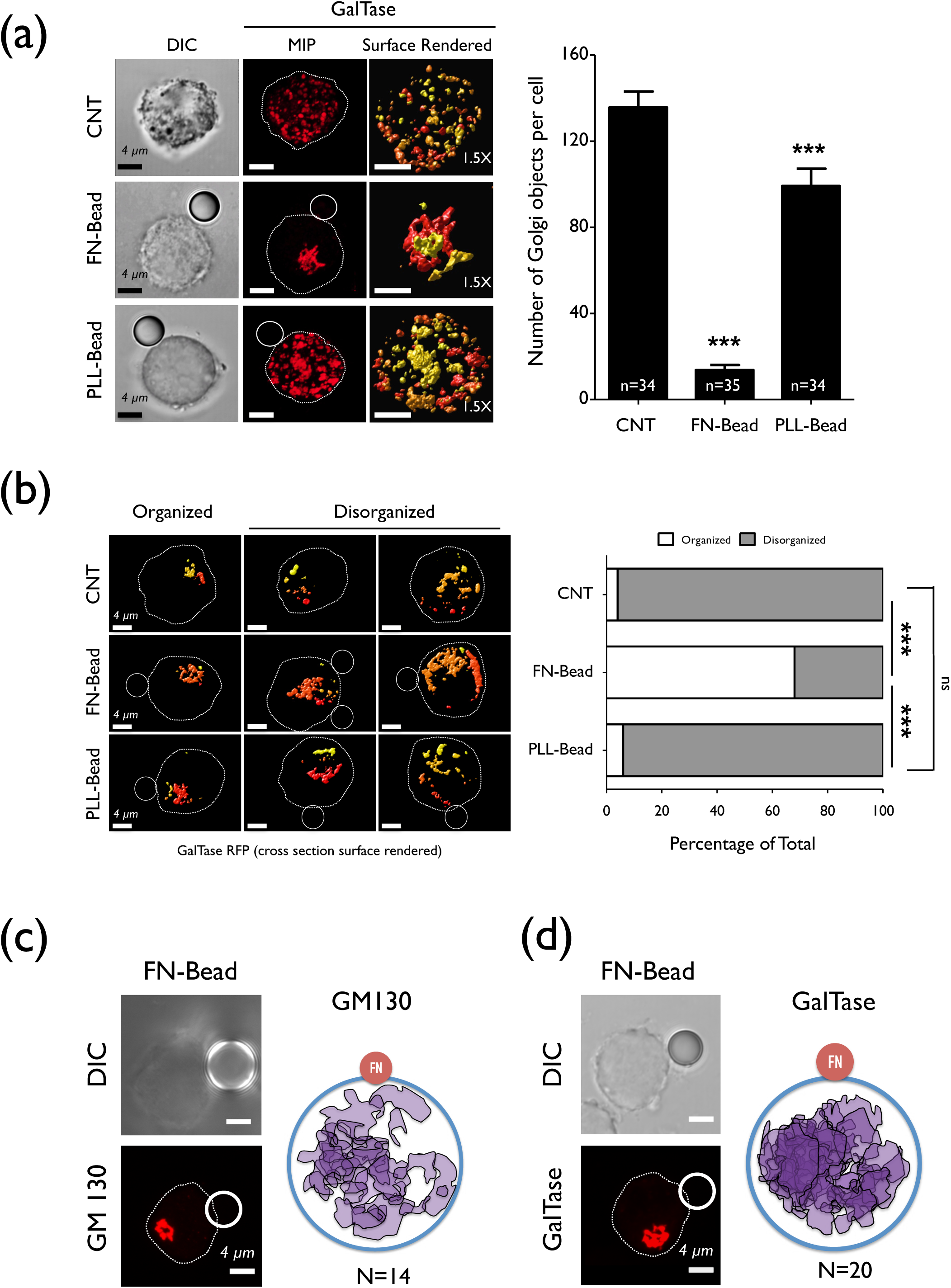
Binding to FN-coated bead restores Golgi organization in suspended cell. **(a)** WT-MEFs expressing GalTase RFP serum starved for 12 hours were detached and held in suspension for 30 mins and incubated without beads (CNT) or with beads coated with fibronectin (FN-Bead) or poly L-Lysine (PLL-Bead) for 15 mins. DIC image of cell shows the attached bead. Their trans-Golgi detected using GalTase RFP (GalTase) was imaged and Z stack images deconvoluted and a representative maximum intensity projection (MIP) image shown. The cell perimeter and bead position are marked here. These images were surface rendered and zoomed for clarity (1.5X). Discontinuous Golgi objects per cell were determined using the Huygens image analysis software. Graph represents mean ± SE from minimum of 34 cells and maximum of 35 cells (as indicated in each bar graph) from three independent experiments. Statistical analysis of this data was done using the Mann Whitney’s test (*** p-value <0.0001). (**b**) The percentage distribution of cells with organized and disorganized Golgi phenotypes in WT-MEFs expressing GalTase RFP incubated without (CNT) or with bead coated with fibronectin (FN-Bead) or poly L-lysine (PLL-Bead) was determined by counting 50 cells at each time point. The left panel shows representative surface rendered cross section images for the organized and disorganized phenotype at each treatment. The graph represents percentage distribution for each phenotype in one of two comparable independent experiments. Statistical analysis comparing the change in distribution profile was done using the Chi-Square test, two-tailed (*** p-value < 0.0001). Scale bar in all the images is set at 4 µm. The position of the (**c**) cis Golgi (GM130) and (**d**) trans Golgi (GalTase) mapped as discussed in methods, relative to fibronectin bead (FN) for 14 and 20 cells respectively (from 3 independent experiments) are represented as a schematic (right panel). The position of the bead from each cell is made to overlap at one spot (represented as a single bead) (FN) with the relative position of the Golgi maintained. A representative DIC image of the cell with the attached bead and the localization of the cis and trans Golgi respectively are shown to the left of each schematic. All scale bars in images are 4 µm.

Fibronectin coated beads bound to cells also offer a means of evaluating the spatial regulation of adhesion-dependent Golgi reorganization. Using cells bound to a single bead the orientation of cis-Golgi (GM130) and trans-Golgi (GalTase) relative to bead were compared across cells. The reorganized Golgi did not show a spatial predisposition for the FN bead (Fig 2c, 2d) in suspended cells or to fibronectin-coated coverslips when re-adherent (Supplementary Fig. 2b, 2c). This does suggest adhesion likely drives reorganization of the Golgi along a pre-existing microtubule network to the MTOC.

### Microtubule network is essential for adhesion-dependent Golgi organization

Knowing the role cytoskeletal networks have in integrin-dependent cellular function (Balasubramanian et al., 2007a; Parsons et al., 2010) and regulation of Golgi architecture (Hurtado et al., 2011; Sandoval et al., 1984; Valderrama et al., 1998), we tested the role it could have in this pathway. Loss of adhesion and re-adhesion did not disrupt the microtubule network (top two panels), MTOC or the actin cytoskeletal network (bottom panel) in cells (Fig. 3a). Both cytoskeletal networks we find to also be functional in non-adherent cells, with their disruption distinctly affecting endocytic trafficking of GM1-CTxB on the loss of adhesion (Fig. 3b). On nocodazole treatment, GM1 is endocytosed into the cell cortex but not trafficked to the recycling endosome (Balasubramanian et al., 2007a) (Fig. 3b middle panel), while Latrunculin A treatment blocks GM1 endocytosis (Fig. 3b right panel). WT-MEFs suspended for 60 mins and then treated with Nocodazole (10 µM) show the cis-medial Golgi (Man II GFP) to be further disorganized with a significant increase in the number of discontinuous Golgi objects detected per cell (Fig. 3e left panel). The trans-Golgi (Galtase RFP), which on the loss of adhesion is already extensively disorganized, did not show any further visible change (Fig. 3e right panel). This suggests the differential distribution of the cis vs trans Golgi in non-adherent cells could be dependent on their relative association with the microtubule network. On re-adhesion, Nocodazole-treated cells failed to reorganize both the cis-medial and trans-Golgi (Fig. 3f). Washout of the nocodazole restores cis-Golgi re-organization in these cells (Supplementary Fig. 3a). Disrupting the actin cytoskeleton with Latrunculin A, however, did not affect the Golgi organization in suspended and re-adherent cells (Fig. 3c, 3d). This suggests the adhesion-dependent regulation of the Golgi to be dependent only on the microtubule network. It also raises the possible role microtubule motor proteins associated with the Golgi could have in mediating the same. The motor protein Dynein helps move Golgi membranes to the microtubule minus end, active GTP bound Arf1 recruiting it to through Golgin 160 to Golgi membranes (Yadav et al., 2012). We hence asked if adhesion could regulate Arf1 activation (like it regulates Arf6) (Balasubramanian et al., 2007a) to control Golgi organization and the role Dynein has in mediating the same.

**Figure 3.**
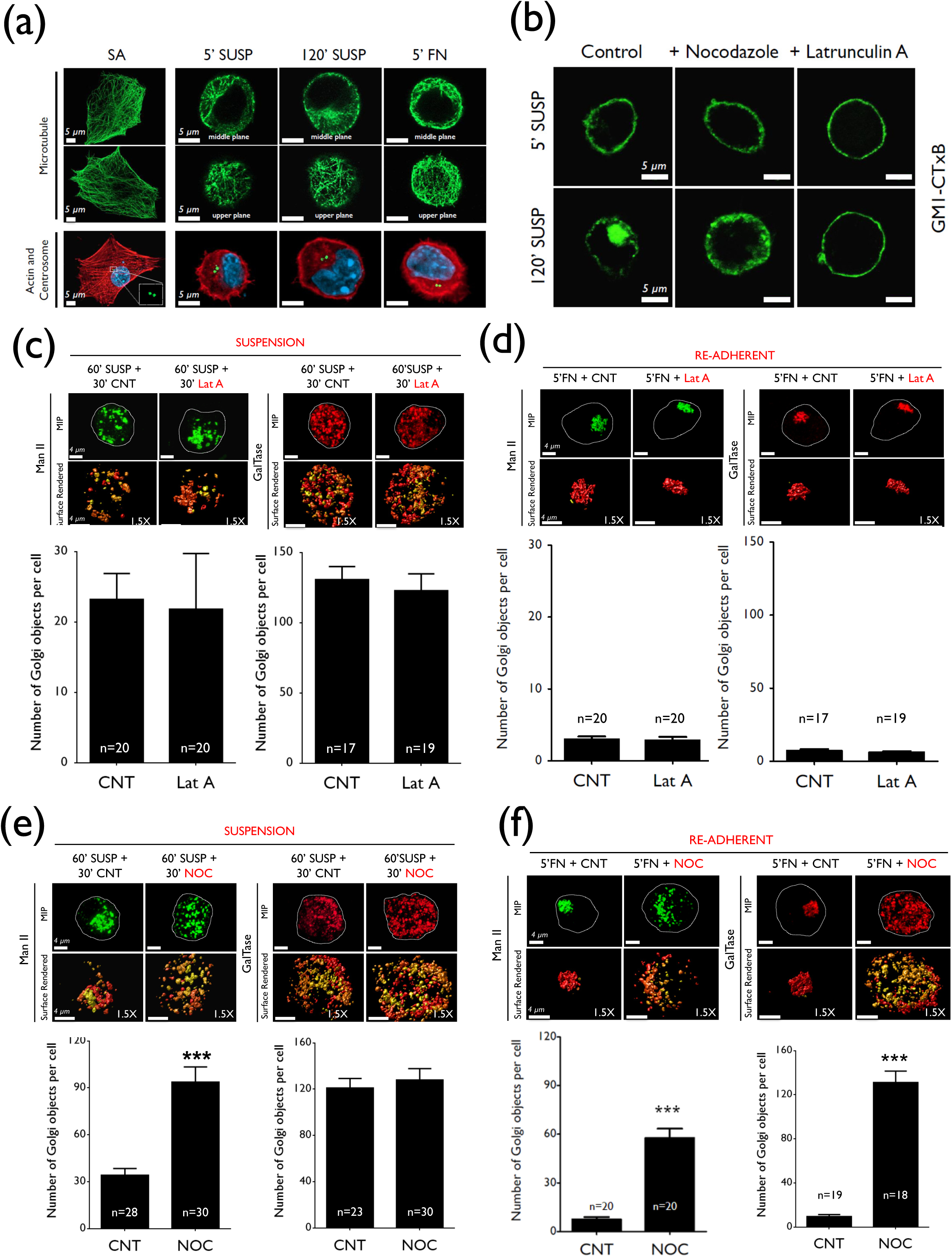
Adhesion-dependent Golgi organization is dependent on the microtubule network. **(a)** Serum-starved stable adherent (SA) WT-MEFs detached (5’ SUSP), held in suspension for 120 mins (120’ SUSP) and re-plated on fibronectin for 5 mins (5’ FN) were immuno-stained for β-tubulin (microtubule). Cross-sectional representative images for a middle (top panel) and upper plane (lower panel) for suspended and re-adherent cells are shown. A middle plane for two representative stable adherent (SA) cells is also shown. Cells stained for actin and γ-tubulin (to detect the centrosome) were deconvoluted and representative images for each time point shown in the lowermost panel. Images are representative of 30 cells imaged from 3 independent experiments. Scale bar in these images is set at 5 µm. (**b**) Stable adherent serum starved WT-MEFs, untreated (control), pre-treated with 10 µM Nocodazole or 0.5 µM Latrunculin A were pre-labeled with cholera toxin B (GM1-CTxB) detached (5’ SUSP) and held in suspension for 120 mins (120’ SUSP). Images are representative of 20 cells (Nocodazole) and 30 cells (Latrunculin A) respectively, from 2 independent experiments. Scale bar in the images is set at 5 µm. Serum-starved WT-MEFs expressing cis-medial (Man II) or trans Golgi (GalTase) marker were suspended for (**c**) 60 mins (60’ SUSP) and an additional 30 mins without (+30 ‘CNT) or with Latrunculin A (+30‘LatA) and (**d**) re-plated on fibronectin for 5 mins without (5’ FN+CNT) or with drug (5’ FN+LatA). Confocal Z stacks were de-convoluted and representative maximum intensity projections (MIP) (top panel) and zoomed (1.5X) surface rendered images are shown (bottom panel). Following deconvolution, discontinuous Golgi objects per cell for Man II and GalTase were determined using the Huygens image analysis software. The graph represents mean ± SE (17–20 cells from two independent experiments). Statistical analysis was done using the Mann Whitney’s test (p-value > 0.05). Scale bar in the images is set at 4 µm. Cells expressing cis Golgi (Man II) or trans Golgi (GalTase) marker held in suspension for (**e**) 60 mins (60’ SUSP) were held in suspension for an additional 30 mins without treatment (+30’ CNT) or treated with Nocodazole (+30’NOC) and (**f**) re-plated on fibronectin for 5 mins without drug (5’FN+CNT) or with nocodazole (5’FN+NOC). Confocal Z stack images were deconvoluted, represented as maximum intensity projections (MIP) (top panel) and surface rendered images zoomed 1.5X for clarity (bottom panel for each cell). Discontinuous Golgi objects per cell for Man II and GalTase were determined using the Huygens image analysis software. The graph represents mean ± SE (18–30 cells from three independent experiments). Statistical analysis comparing CNT and NOC were done using the Mann Whitney’s test (*** p-value <0.001). Scale bar in the images is set at 4 µm.

### Adhesion dependent Arf1 activation regulates Golgi organization

Active Arf1 pulled down using GST conjugated GGA3 effector binding domain reveals loss of adhesion to significantly reduce Arf1 activation (by ~60%), rapidly restored on re-adhesion to fibronectin (Fig. 4a). Net Arf1 levels do not change under these conditions (Supplementary Fig. 4a). To test if this drop in active Arf1 levels has a role in mediating Golgi disorganization we disrupted the same by expressing GFP tagged constitutively active Arf1 (Q71L) in WT-MEFs (Supplementary Fig. 4b) and find it prevents disorganization of the trans-Golgi (Galtase-RFP) in suspended cells (Fig. 4b). GFP WT Arf1 expressing MEFs behave like control cells, with a distinctly disorganized Golgi phenotype (Fig. 4b). This is further reflected in a significant decrease in the number of trans Golgi objects per cell in suspended Q71L Arf1 expressing cells relative to WT Arf1 and control (Graph in Fig. 4b). The distribution profile of organized versus disorganized phenotype in suspended and re-adherent WT Arf1 and Q71L Arf1 expressing cell populations further confirm this regulation (Supplementary Fig. 4c). Similar expression of HA-tagged active Q71L Arf1 (Supplementary Fig. 4f) in suspended cells was found to block cis-Golgi (GM130) disorganization (Supplementary Fig. 4g). Constitutively active Arf6 (T157A) similarly expressed in cells did not have this effect (Supplementary Fig. 4i) further confirming this regulation to be Arf1 specific.

**Figure 4.**
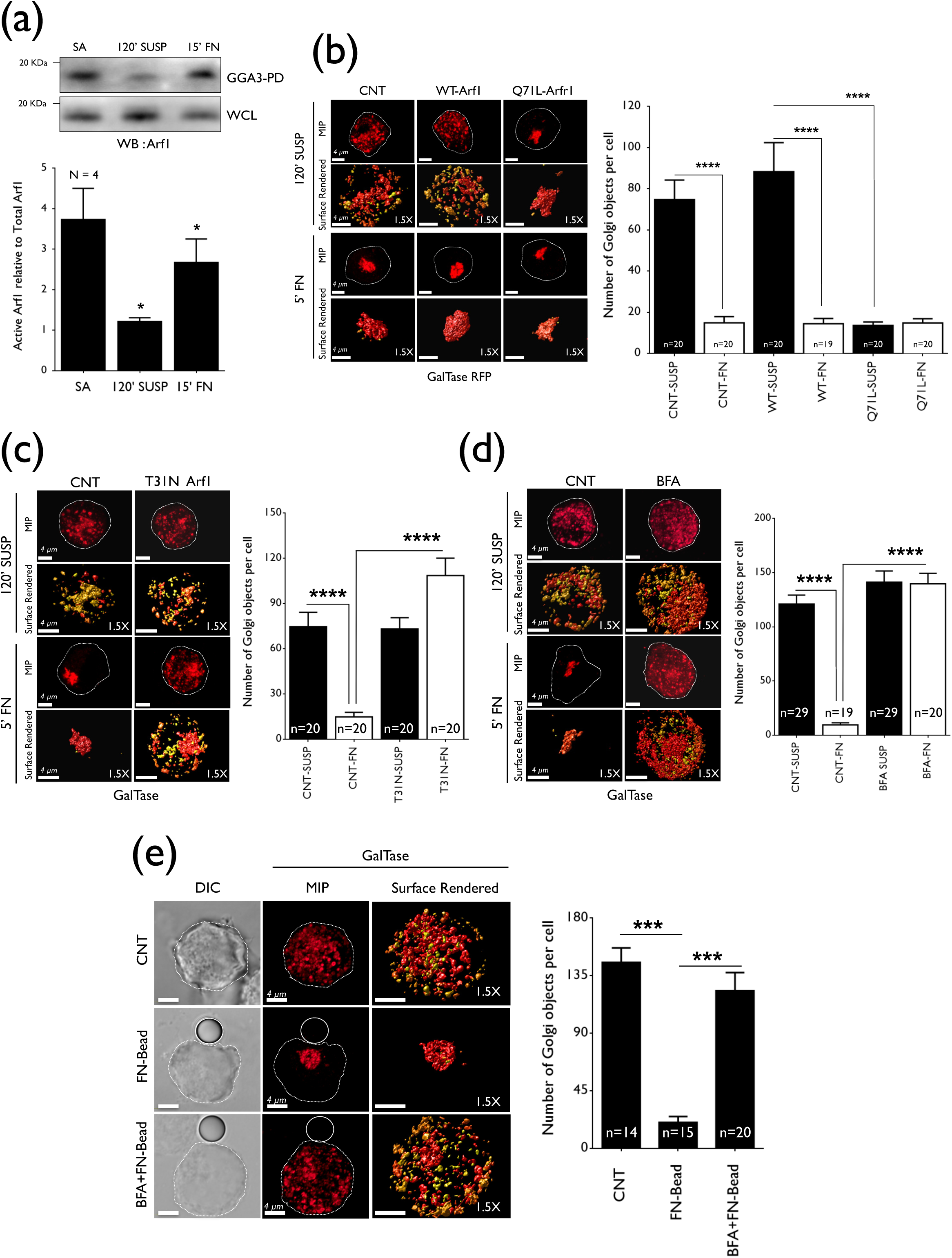
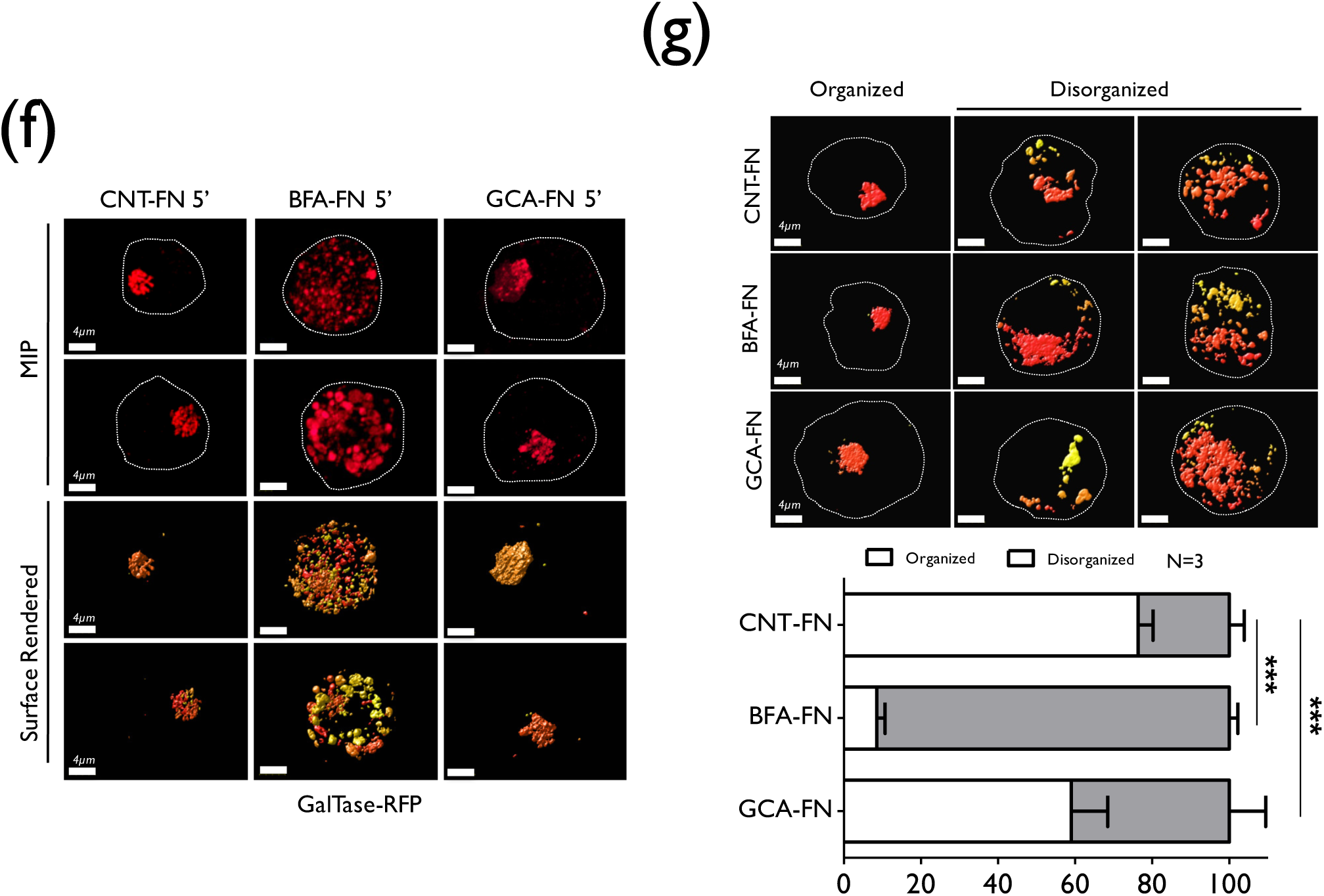
Adhesion-dependent Arf1 activation regulates Golgi organization. **(a)** Western blot detection of active Arf1 in GST-GGA3 pulldown (GGA3 PD) and total Arf1 in whole cell lysate (WCL) of serum-starved WT-MEFs stable adherent (SA), suspended for 120 mins (120’ SUSP) and re-adherent on fibronectin for 15 mins (15’ FN). The graph represents mean ± SE from four independent experiments. (**b**) Serum-starved WT-MEFs transfected with GalTase RFP alone (CNT) or along with GFP wild-type Arf1 (WT-Arf1), active Arf1 (Q71L Arf1) or (**c**) dominant negative Arf1 (T31 N Arf1) were detached, held in suspension for 120 mins (120’ SUSP) and re-plated on fibronectin for 5 mins (5’FN). (**d**) Cells expressing GalTase RFP were held in suspension for 90 mins and treated with MeOH (CNT) or brefeldin A (BFA) for additional 30 mins (120’ SUSP) and re-plated on fibronectin for 5 mins (5’FN) without (CNT) or with BFA (BFA). (**e**) Control and BFA treated suspended cells incubated with beads coated with fibronectin (FN-Bead) and (BFA+FN-Bead) for 15 mins or MeOH as mock (CNT). In all of the above confocal Z stacks were deconvoluted and representative maximum intensity projection (MIP) and surface rendered image zoomed 1.5X is shown. Discontinuous Golgi objects per cell determined using Huygens is represented in the graph as mean ± SE from a minimum of 15 and maximum of 29 cells from two (**b, c**, and **e**) and three (d) independent experiments. (**f**) Cells expressing GalTase RFP held in suspension for 90 mins and treated with DMSO (mock) brefeldin A (BFA, 10 µg/ml) or Golgicide-A (GCA, 10 µM in DMSO) for additional 30 mins (120’ SUSP) were re-plated on fibronectin for 5 mins (5’FN) without drug (CNT FN-5’) or with BFA (BFA FN-5’) or with GCA (GCA FN-5’). (**g**). Percentage distribution of cells with organized and disorganized Golgi phenotypes determined and representative surface rendered cross section images shown. The graph represents mean ± SE from 3 independent experiments. Statistical analysis was done using the two-tailed Chi-Square test (*** p-value <0.0001). All scale bars in images are set at 4 µm.

We further tested if re-adhesion mediated recovery of Arf1 activation is needed for Golgi reorganization, by inhibiting Arf1 using a dominant negative mutant (Donaldson and Jackson, 2000) (GFP tagged T31 N Arf1) (Supplementary Fig. 4b) or inhibitor Brefeldin A (BFA) (Fujiwara et al., 1988; Lippincott-Schwartz et al., 1989). Cells suspended for 90 mins were incubated with BFA for an additional 30 mins and then re-plated on fibronectin. Both treatments while not further disrupting the disorganized trans-Golgi phenotype (GalTase-RFP) in suspended cells (120’ SUSP) did prevent the Golgi from reorganizing on re-adhesion to fibronectin (Fig. 4c, 4d). This is further reflected in Golgi object numbers in re-adherent cells being comparable to those seen in suspended cells (Graphs in Fig. 4c, 4d). Similar expression of HA-tagged dominant negative T31 N Arf1 (Supplementary Fig. 4f) in suspended cells also prevented cis-Golgi (GM130) from reorganizing in re-adherent cells (Supplementary Fig. 4h). The distribution profile of organized versus disorganized phenotype in BFA treated suspended and re-adherent cell populations confirming the same (Supplementary Fig. 4d). Binding of fibronectin-coated beads to suspended BFA treated cells (BFA+FN-Bead) failed to restore Golgi integrity unlike untreated cells bound to FN beads (Fig. 4e). Golgi object numbers accordingly stayed significantly high in treated cells (Graph in Fig. 4e). The distribution profile of the organized versus disorganized phenotype in BFA treated cells further confirming this (Supplementary Fig. 4e).

BFA acts as a non-competitive inhibitor of Arf GEFs, BIG1/2 (BFA-inhibited GEFs) and GBF1 (Golgi-specific Brefeldin A-resistance factor 1) (Casanova, 2007). We further compared the effect BFA has to Golgicide A, known to specifically target only GBF1 (Sáenz et al., 2009). While BFA blocked Golgi reorganization dramatically Golgicide A only had a minor effect on the same, reflected in their distribution profiles (Fig. 4f, 4g). This suggests BIG1/2 (over GBF1) is the prominent GEF working downstream of adhesion to regulate Arf1 activation and Golgi organization.

### Adhesion-dependent Arf1 activation regulates its binding to dynein

Activation of Arf1 is seen to be dependent on its localization at the Golgi, which could, in turn, mediate recruitment of the minus end motor protein dynein (Yadav et al., 2012). We hence tested if differential Arf1 activation by adhesion could regulate its binding and recruitment of the dynein in suspended vs re-adherent cells driving the observed Golgi phenotype. Pull down of active Arf1 with GST-GGA3 we find does bring down dynein with it, inhibition of Arf1 with BFA affecting the same (Supplementary Fig. 5a). Loss of adhesion mediated drop in active Arf1 (Fig. 5a left panel) also significantly reduces the amount of dynein that is brought down with it, re-adhesion restoring active Arf1 and bound dynein levels (Fig. 5a right panel). This suggests adhesion-dependent Arf1 activation can differentially recruit dynein to drive Golgi reorganization. Accordingly, ciliobrevin mediated inhibition of dynein blocks re-adhesion mediated Golgi organization (Fig. 5b, Supplementary Fig. 5b) reflected in Golgi object numbers (Graph in Fig. 5b). Ciliobrevin did not affect adhesion-dependent Arf1 activation (relative to untreated control) (Fig. 5c) or Arf1 levels (Supplementary Fig. 5c), suggesting the inhibition of dynein downstream of Arf1 is what affects Golgi reorganization here. Together this reveals the presence of an adhesion-Arf1-Dynein-Microtubule pathway mediating Golgi organization in cells.

**Figure 5.**
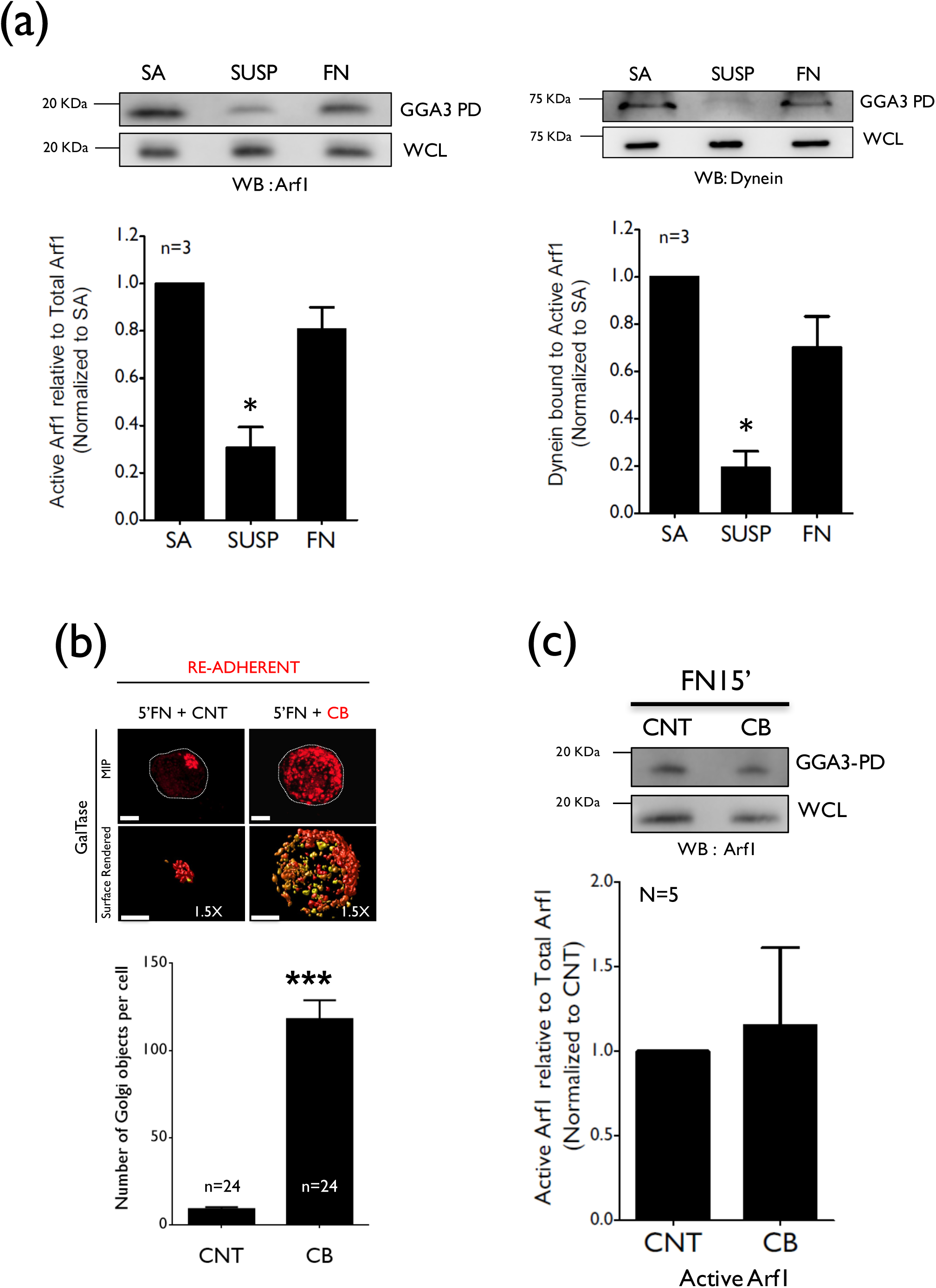
Adhesion-dependent activation of Arf1 recruits dynein for Golgi organization. **(a)** Western blot detection of active Arf1 (left panel) and dynein (right panel) in GST-GGA3 pulldown (GGA3 PD) and total Arf1 & dynein in whole cell lysate (WCL) of serum-starved WT-MEFs stable adherent (SA), suspended for 120 mins (120’ SUSP) and re-adherent on fibronectin for 15 mins (15’ FN). The ratio of densitometric band intensities of Arf1 and dynein in pulldown relative to their levels in the WCL are represented in the graph as mean ± SE from four independent experiments. (**b**) Cells expressing GalTase RFP were held in suspension for 90 mins (CNT) and treated with Ciliobrevin-D (CB, 20 µM in DMSO) for additional 30 mins and re-plated on fibronectin for 5 mins without (5’FN+CNT) or with CB (5’FN+CB). Confocal Z stacks deconvoluted and representative maximum intensity projection (MIP) and surface rendered zoomed image (1.5X) shown. Discontinuous Golgi objects per cell determined using Huygens represented in the graph. Mean ± SE from three independent experiments. Statistical analysis comparing object numbers was done using the Mann Whitney’s test (*** p-value <0.0001). All scale bars in images are set at 4 µm. (**c**) Western blot detection of active Arf1 using GST-GGA3 pulldown (GGA3 PD) and total Arf1 in whole cell lysate (WCL) of serum-starved WT-MEFs suspended for 120 mins and re-adherent on fibronectin for 15 mins (FN 15’) without (CNT) or with ciliobrevin (CB). The ratio of densitometric band intensities of Arf1 in the pulldown relative to their levels in the WCL is represented in the graph as mean ± SE from five independent experiments.

### Adhesion-dependent Golgi organization affects Golgi function

We further asked if and how this pathway regulates Golgi function. A major read out of Golgi function in cells is their ability to glycosylate and deliver protein and lipids at the plasma membrane. Both *N*- and *O*-glycosylation involve a series of enzymatic reactions catalyzed by glycan-processing enzymes across the cis, medial and trans-Golgi compartments (Stanley, 2011; Varki, 1998). Changes in Golgi architecture does affect processing and trafficking of glycosylated proteins and lipids (Pokrovskaya et al., 2011) and can be detected using lectins that selectively recognize glycan epitopes (Sharon and Lis, 2004). Using flow cytometry, we quantitated the cell membrane binding of fluorescently tagged lectins, Concanavalin A (ConA) (mannose-binding), wheat germ agglutinin (WGA) (Galactose/N-acetylgalactosamine binding), peanut agglutinin (PNA) (N-acetylglucosamine binding) and ulex europaeus agglutinin (UEA) (Fucose binding), on loss of adhesion. Cells held in suspension for 120 mins show a significant increase in plasma membrane binding of all four lectins (black bars) relative to their basal levels when detached (grey bars) (Fig. 6a). This suggests the loss of adhesion promotes Golgi processing and/or trafficking to increase membrane glycosylation levels. To confirm if this is indeed the result of the disorganized Golgi phenotype on the loss of adhesion we tested if active Arf1 mediated restoration of Golgi integrity in suspended cells could prevent this glycosylation change. Cherry tagged WT Arf1 and active Q71L Arf1 (Supplementary Fig. 6a) expressing cells show comparable Arf1 expression (Supplementary Fig. 6b) and only a modest change in basal cell surface ConA and UEA binding when detached (5’ SUSP) (Supplementary Fig. 6c, 6d). Active Arf1 did, however, block the increase in cell surface glycosylation observed in suspended control and WT Arf1 expressing cells (Fig. 6b, 6c). This confirms adhesion-dependent regulation of Arf1 and resulting Golgi disorganization affects processing, making it a vital regulator of Golgi function in cells.

**Figure 6.**
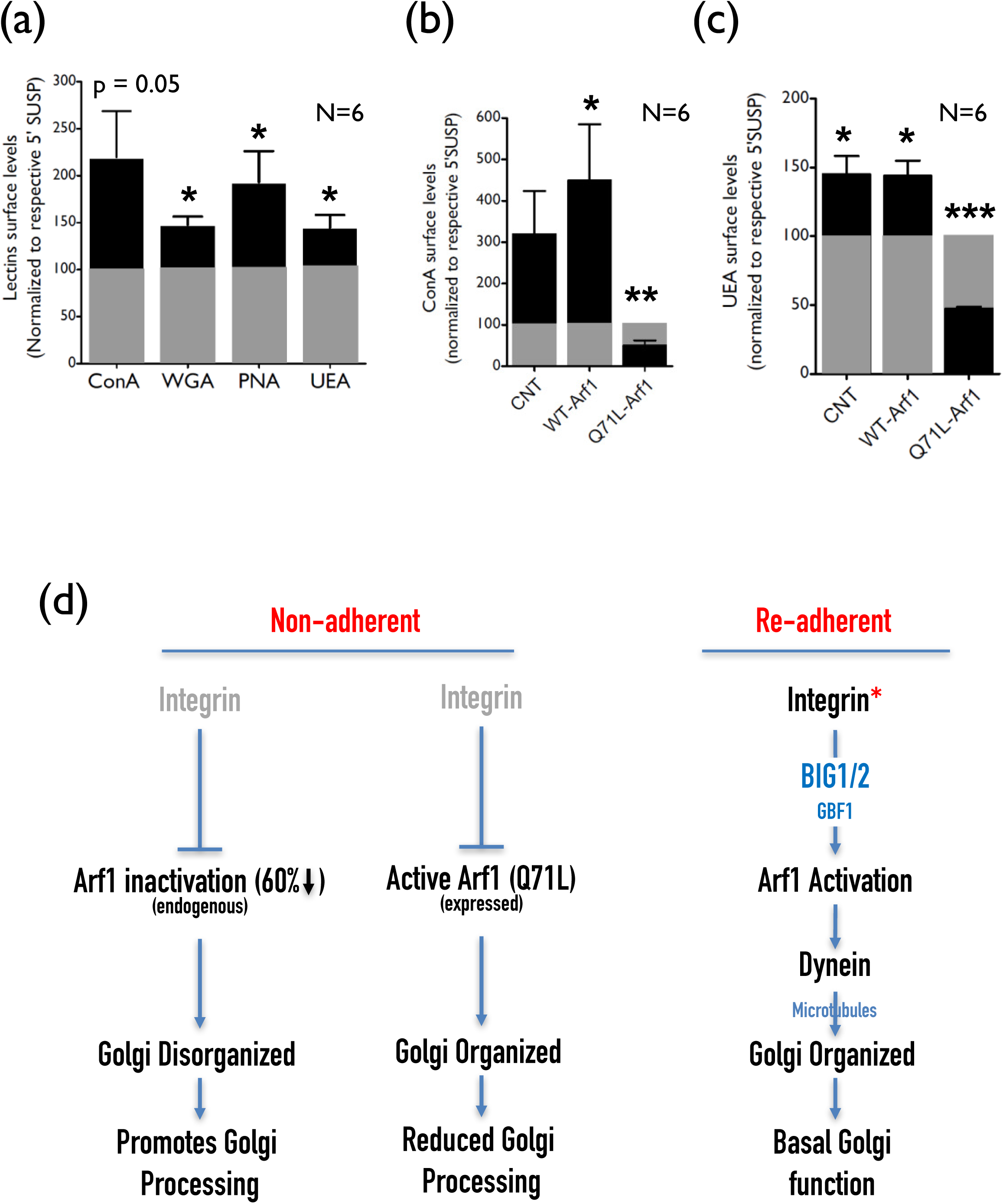
Loss of adhesion mediated Golgi disorganization affects Golgi function. **(a)** Serum-starved WT-MEFs detached with Accutase (5’ SUSP) and held in suspension for 120 mins (120’ SUSP), labeled with Alexa-488 conjugated ConA, WGA, PNA and FITC-UEA lectin. Cell surface-bound lectin fluorescence was measured by FACS and median fluorescence intensity of 120’ SUSP cells (black bar) was normalized to their respective 5’ SUSP cells (equated to 100 and represented as a grey bar). The graph represents mean ± SE from 6 independent experiments. WT-MEFs expressing cherry-N1 (CNT), wild-type Arf1-Cherry-N1 (WT-Arf1) or active Arf1-Cherry-N1 (Q71L Arf1) were similarly labeled with (**b**) ConA-Alexa488 and (**c**) UEA-FITC. Surface-bound lectin fluorescence in cell population gated for cherry tagged Arf1 expression was measured by FACS and median fluorescence intensity at 120’ SUSP (black bar) normalized to their respective 5’ SUSP cells (equated to 100 and represented as a grey bar). The graph represents mean ± SE of 6 independent experiments. (**d**) Schematic represents the integrin-dependent regulation of Arf1 through BIG1/2, that allows for the recruitment of dynein which along the microtubule network regulates Golgi architecture and function.

### Discussion

This study hence reveals cell-matrix adhesion to be a novel regulator of Golgi architecture and function (Fig. 6d). Independent of growth factors, loss of adhesion rapidly disorganizes the Golgi, restored on re-adhesion to fibronectin for 5 mins or less. Such early adhesion events in serum-deprived cells are likely mediated through integrins. The restoration of integrin-dependent Akt activation (del Pozo et al., 2004a) in these cells supports the same. This is further established by the fact that in these cells binding to fibronectin-coated beads (but not poly-L-lysine), seen to cause integrin clustering and activation (del Pozo et al., 2004a), completely reorganizes the Golgi. The effect limited integrin activation at the site of bead attachment has on the Golgi is interestingly not spatially restricted. This not only suggests small changes in integrin activation and clustering is sufficient to drive this pathway, but also raises the possibility that the signal downstream of integrins could indeed dissipate very rapidly to the Golgi. One such regulator working downstream of integrins could be calcium. Studies have shown integrin activation can drive transient changes in intracellular Ca^+2^ levels (Sjaastad et al., 1996). Studies in neuronal cells have shown a calcium-dependent interaction between NCS-1 and ARF1 that can control Golgi architecture. This suggests a functional cross-talk could exist between Ca^2+^-dependent and ARF-dependent pathways in the trans-Golgi (Haynes et al., 2005). Our study for the first time identifies adhesion to control Arf1 activation, regulated by the BFA sensitive Arf1 GEF BIG1/2.

Earlier studies have suggested BIG1/2 and GBF1 (both sensitive to BFA) to differentially localize in trans and cis Golgi respectively (Manolea et al., 2008). Thus, on the loss of adhesion BIG1/2 mediated Arf1 activation is prominently downregulated in the trans-Golgi, relative to the cis/cis-medial Golgi (containing GBF1). This could reflect in the greater disorganization of the trans-Golgi in non-adherent cells. While such a differential role and regulation of trans vs cis/cis-medial is speculated (Rothman, 1982; Sens and Rao, 2013), loss of adhesion is among the few physiological processes that visibly affect their organization differently. While net Arf1 activation on the loss of adhesion drops by ~60%, the effect this drop has on local active Arf1 levels in the trans vs cis/cis-medial Golgi could indeed vary and remains to be confirmed. This could, in turn, affect the ability of active Arf1 to recruit dynein, to differentially affect their organization. A role for plus-end motor protein, kinesin, in this pathway also remains to be explored. Kinesin, KIF1C, is seen to bind the Golgi protein Rab6A directly, regulating its binding to the microtubule network, to regulate Golgi fragmentation (Lee et al., 2015). It is hence possible that dynein and kinesin motors together control Golgi membrane movement on microtubules to control their organization by adhesion.

The sensitive, rapid and reversible nature of this regulation suggests cells could use this pathway to fine-tune Golgi function. A change in cell surface glycosylation by adhesion-dependent regulation of Golgi architecture could indeed affect multiple aspects of cellular function (Stanley, 2011). Most cell surface receptors are glycosylated through the Golgi (Apweiler et al., 1999), with it known to affect their conformation(Takahashi et al., 2009), ligand binding capability (Bellis, 2004), receptor dimerization(Sato et al., 2009) and downstream signalling(Zhao et al., 2008). Glycosylation of β1 integrin (β1) is seen to regulate its function in matrix adhesion, migration (Isaji et al., 2004) and seen to be regulated by BIG1 mediated Arf1 activation (Shen et al., 2007). Malignant transformation in cancer cells is also accompanied by aberrant glycosylation of proteins, including integrins and cadherins (Bassagañas et al., 2014; Kariya et al., 2017). In many cancers, the Golgi is fragmented to drive this change (Migita and Inoue, 2012; Petrosyan, 2015). It will be of much interest to test if and how changes in surface glycosylation on the loss of adhesion compare to those seen on oncogenic transformation and how they contribute to anchorage-independent signaling (Guo and Giancotti, 2004; Schwartz, 1997). Integrin-mediated adhesion is seen to regulate endocytic and exocytic trafficking, to control anchorage-dependent signaling, that is deregulated in cancers (Balasubramanian et al., 2007a, 2010; Pawar et al., 2016). The Golgi is a vital regulator of protein sorting and trafficking(Hwang, 2008) and adhesion by regulating its organization could further control the same.

While the loss of adhesion and re-adhesion are rare occurrences for most anchorage-dependent cells the sensitive nature of this regulation does suggest that changes in the extent of adhesion could trigger a reorganization of the Golgi to affect its function. Integrin-mediated adhesion is known to regulate cell cycle progression, helping determine spindle orientation and axis in a dividing cell (Lancaster et al., 2013). Golgi fragmentation and partitioning are necessary for cells to progress through mitosis (Rabouille and Kondylis, 2007). This is mediated by the regulation of Arf1. The recruitment of Arf1 to Golgi membranes is inhibited early in mitosis by its inactivation. Arf1-dependent proteins fail to be recruited from the cytoplasm to Golgi membrane causing it to fragment and fall back into the ER (Altan-Bonnet et al., 2003). Further, expressing Arf1[Q71L] was seen to prevent this Golgi fragmentation in mitosis (Altan-Bonnet et al., 2003). This mitotic Golgi fragmentation is associated with mitotic cell rounding that involves a change in cell shape and adhesion (Cadart et al., 2014; Dao et al., 2009). The disorganized Golgi phenotype on loss of adhesion we find is distinctly different from a BFA mediated Golgi fragmentation, that causes the cis-Golgi to rapidly disorganize and fall back into the ER (Fujiwara et al., 1988; Lippincott-Schwartz et al., 1989). In non-adherent cells the disorganized cis-Golgi shows almost no overlap with the ER (Fig. 1g, 1h), till these cells are treated with BFA. While in non-adherent cells a 60% decrease in active Arf1 levels (relative to stable adherent cells) is observed, BFA causes an additional 75% reduction in Arf1 activation that drives its fall back into the ER (data now shown). This could be mediated by BFA mediated inhibition of GBF1 in the cis/cis-medial Golgi in a way that loss of adhesion does not. This suggests, changing Arf1 inactivation could progressively affect Golgi organization, making the disorganized Golgi seen on loss of adhesion a possible structural (and maybe functional) intermediate to Golgi fragmentation. This study hence not only identifies cell-matrix adhesion as a novel regulator of Golgi structure and function, but in doing so highlights the possible role such a regulatory pathway could have in normal and disease conditions.

## ACKNOWLEDGEMENTS

Flow Cytometry studies were supported by Sujaya Ingale, at the DBT-BIRAC supported Venture Center BioIncubator at CSIR-NCL, Pune, India. We thank Dr. Jennifer Lippincott-Schwartz (NICHD, NIH, Bethesda) for generously providing GalTase-RFP, ManII-GFP and KDEL-REP constructs. We thank Dr. Satyajit mayor (NCBS, Bangalore, India) for generously providing WT and T31 N-Arf1 constructs. We thank Remko Dijkstra at SVI for help with image analysis using the Huygens professional software. We thank Martin Schwartz for discussions during the course of the study. We thank IISER Pune for the facilities and infrastructure. VS thank Infosys Foundation Travel Award.

## MATERIALS AND METHODS

### Reagents

Fibronectin was purchased from Sigma (Cat. No. # F2006). Cholera toxin subunit B (CTxB) conjugated with Alexa 594 (C22843) or Alexa 488 (C34775) were purchased from Invitrogen Molecular Probes and used at a 1:10000 dilution. Accutase was purchased from Sigma (Cat. No. # A6964). Lectin probes were purchased Invitrogen Molecular Probes, Concanavalin A-Alexa 488 (Cat. No. # C11252), PNA-Alexa-488 (Cat. No. # L21409), WGA-Alexa-488 (Cat. No. # W11261). UEA-FITC was purchased from Sigma (Cat. No. # L9006). Divinyl Polystyrene beads (Cat. No. # 42045A 1) were purchased from Thermo scientific. Nocodazole (Cat. No. # M1404), Latrunculin A (Cat. No. # L5163), Brefeldin A (Cat. No. # B7651), Golgicide A (Cat. No. # G0923) were purchased from Sigma. Ciliobrevin D (Cat. No. # 250401) was purchased from Calbiochem. Fluoromount-G (Cat. No. # 0100–01) was purchased from Southern Biotech.

### Antibodies

Antibodies used for western blotting include mouse anti-Arf1 (clone 1D9, abcam, Cat. No. # ab2806) at a dilution of 1:500, Rabbit anti-Arf1 (clone EP442Y, abcam, Cat. No. # ab32524) at a dilution of 1:500, Mouse anti Dynein (clone 74.1, Millipore, Cat. No. # MAB1618) at a dilution of 1:2000, rabbit anti GFP (Santa Cruz, Cat. No. # GFP (FL): sc-8334) at a dilution of 1:700, Mouse anti βeta-tubulin (Clone E7, Developmental Studies Hybridoma Bank, Cat. No. #AB_2315513) at a dilution of 1:5000, mouse anti-HA.11 Epitope Tag Antibody (Clone 16B12, Covance, Cat. No. # MMS-101 R) at a dilution of 1:2000, Secondary antibodies conjugated with HRP were purchased from Jackson Immunoresearch, and were used at a dilution of 1:10000.

Antibodies used for immunofluorescence in this study include mouse anti-GM130 (BD transduction, clone 35, Cat. No. # 610822) at a dilution of 1:100, Mouse anti βeta-tubulin (Clone E7, Developmental Studies Hybridoma Bank, Cat. No. #AB_2315513) at a dilution of 1:1000, Anti-gamma Tubulin antibody (Abcam, Cat. No. # ab11317) at a dilution of 1:100, Rat anti-HA Epitope Tag Antibody (Clone 3F10, Roche, Cat. No. # 11867423001) at a dilution of 1:1000. Alexa 594 Phalloidin (Invitrogen ThermoFisher Scientific, Cat. No. # A12381) was used at a dilution of 1:100. Secondary antibodies with Alexa conjugate (488 or 594) were purchased from Invitrogen Molecular Probes (Cat. No. # A12379 and A12381) and were used at a dilution of 1:1000.

### Plasmids

GFP-tagged Arf1-WT and Arf1-T31 N constructs were obtained from Dr. Satyajit Mayor (National Centre for Biological Sciences, Bangalore, India). GFP-tagged Arf1-Q71L construct was made by site-directed mutagenesis using GFP-Arf1-WT as the template and following primers – (forward)5’-GACGTGGGTGGCCTGGACAAGATCCGG-3’ and (reverse) 5’-CCGGATCTTGTCCAGGCCACCCACGTC-3’. mCherry-tagged Arf1-WT and Arf1-Q71L constructs were made by releasing the Arf1 gene from GFP constructs (using Bgl II and BamH1 sites) and cloning the same into an empty mCherry-N1 vector. Galtase-RFP, MannosidaseII-GFP, and KDEL-RFP constructs were all obtained from Dr. Jennifer Lippincott Schwartz (NIH). All of the above-mentioned constructs were sequenced to confirm their identity before being used in our studies.

### Cell culture and transfections

Mouse embryonic fibroblasts (MEFs) obtained from Dr. Richard Anderson (University of Texas Health Sciences Centre, Dallas TX) were cultured in complete Dulbecco’s modified Eagle’s medium (DMEM) (Invitrogen) with 5% fetal bovine serum (FBS) and penicillin-streptomycin (Invitrogen) at 37°C in a 5% CO_2_ incubator. Human foreskin fibroblasts (BJ) Cells from ATCC (ATCC CRL-2522) and were cultured in complete Dulbecco’s modified Eagle’s medium (DMEM) (Invitrogen) with 10% fetal bovine serum (FBS) and penicillin-streptomycin (Invitrogen). WT-MEFs and BJ cells were transfected using LTX-PLUS (Invitrogen) according to the manufacturer’s protocol. Transfections were done in 6 cm dish using 4 µg DNA for 12 hours. 36 hours after transfection cells were serum starved for 12hours with DMEM containing 0.2% FBS.

### Suspension and re-adhesion of cells

WT-MEFS or Human fibroblasts were all cultured in their respective growth medium in 60 mm dishes to ~ 75% confluency. Cells were serum starved (incubated with medium containing 0.2% FBS) for 12 hours detached using Trypsin-EDTA (Invitrogen) or Accutase (Sigma) (for lectin labeling experiments), washed with 0.2% FBS containing DMEM, processed on the ice and an aliquot of cells collected. This processing takes about 5 mins and these detached cells are accordingly labeled as (detached 5’). Cells were then held in suspension with 1% methylcellulose containing low serum DMEM (0.2% FBS) for the required time. Post incubation cells were collected at the required time, carefully washed twice with 0.2% FBS DMEM at 4°C to avoid clumping. These processed cells when required were re-plated on coverslips coated with 2 µg/ml fibronectin for the required time (5 mins, 15mins or 4 hours for stable adherent cells). For confocal microscopy suspended or re-adherent cells were fixed with 3.5% paraformaldehyde (PFA) for 15 mins at room temperature (RT), washed with PBS thrice, stained and mounted using fluoramount. For western blotting, suspended and re-adherent cells were lysed in the required amount of 1X Lamelli, heated at 95°C for 5 mins and stored at -80°C.

### GM1 labeling of cells with CTxB

Serum-starved WT-MEFs detached with trypsin were labeled for 15 mins on ice with cholera toxin B subunit (CTxB) conjugated to Alexa-488 or Alexa-594. These cells were washed and fixed to look at surface GM1 labelling, to begin with, held in suspension for 120 mins and endocytosis of GM1-CTxB detected. Cells were fixed with 3.5% PFA, mounted and observed. To test the role of the cytoskeleton in GM1 endocytosis adherent cells pre-treated with Nocodazole (10 µM) or Latrunculin A (0.5 µM) for one hour were detached, labeled as above and GM1-CTxB endocytosis similarly observed.

### Measuring Cell volume using surface GM1 labeled suspended and re-adherent cells

Z stacks of surface GM1 labeled cells were deconvoluted and surface rendered to determine cell shape in suspended and re-adherent cells. A threshold of 2% was set in order to completely fill the cell and hence giving a continuous structure. The average volume for suspended and re-adherent cells was then compared.

### Binding of fibronectin and poly L-Lysine coated beads to cells

8×10^8^ Divinyl polystyrene beads were re-suspended in 500 µl PBS and sonicated for 30 seconds. Beads were washed thrice with cold 1X PBS (pH 7.4), centrifuged at 6000 rpm for 5 mins at 4°C and incubated with 500 µl PBS containing 10 µg/ml Fibronectin or 10 µg/ml Poly-L-Lysine at 4°C overnight on a rotary shaker. Coated beads were centrifuged, washed with cold 1X PBS (pH 7.4) and blocked with 50 mg/ml BSA at 4°C for 3 hrs on a rotary shaker. Beads were washed and re-suspended finally in 100 µL of PBS (pH7.4). Adherent WT-MEFs were labeled with CTxB-Alexa-594 on ice for 15 mins and then washed thoroughly to remove unbound probe. FN-coated beads (cells to bead ratio 1:10) were added and incubated for 15 mins at 37°C. Cells were then PFA fixed and mounted using Flouromount-G. Cells with bound bead were then imaged (del Pozo et al., 2004b).

Untransfected or GalTase RFP transfected serum-starved cells (2×10^5^) were detached, held in suspension for 30 mins in 2 ml 0.2% FBS containing DMEM with 1% methylcellulose. 12.5 µl FN or PLL coated bead suspension (2×10^6^ beads) was added to the cells (cells: bead ration maintained at 1:10) mixed gently and incubated for 15 mins at 37°C. Cells were washed gently and fixed with 3.5% PFA for 15 mins and RT. Immunostaining of cells for GM130 was done as discussed later, with all washing steps done very carefully. Cells were then mounted using Fluoromount-G, allowed to dry for 24 hours and then imaged. Cells with bound beads were identified in the population and used for comparison.

### Spatial Golgi localization relative to the cell-bound fibronectin coated bead

Cross-sectional images of cells with a single attached FN bead were selected and an outline of the cell boundary with the location and outline of the bead made. The Golgi area was then mapped within the cell. These line drawings for cell outline + bead outline + mapped Golgi area for each cell were grouped together and the position of beads across cells was moved to cause them to overlap. The relative position of the Golgi for each cell while maintained relative to the bead was now comparable across cells and for bead position. This now allows us to generate a combined image for all cells with their cell perimeter and bead position being identical and lets us look at the Golgi position relative to the same. Golgi outlines are made transparent to allow us to see the overlap in localization.

### Inhibitors studies in suspended and re-adherent cells

For all the inhibitor studies, cells transiently expressing GalTase-RFP and ManII-GFP were serum starved with 0.2% FBS containing DMEM for 12 hours, trypsinized and held in suspension for 60 mins in 5ml of 1% methylcellulose containing DMEM. Cells were then treated with Nocodazole (10 µM in DMSO), Latrunculin A (0.5 µM in DMSO), BFA (10 µg in MeOH), Golgicide A (10 µM in DMSO) or Ciliobrevin D (20 µM in DMSO) and incubated for an additional 30 mins at 37°C. Control cells were treated with an equivalent volume of solvent (DMSO /MeOH). Cells were processed as described in suspension assay and samples collected at required times. Cells were re-plated on FN-coated coverslips with or without inhibitor for 5 mins, fixed, mounted and imaged using a confocal microscope.

### Inhibitor studies: Nocodazole washout assay

Serum-starved WT-MEFs were detached and held in suspension for 60 mins and then incubated with Nocodazole (10 µM in DMSO) for 30 mins. Suspended cells were washed twice with PBS to remove methylcellulose in presence or absence of Nocodazole and suspension aliquots fixed with 3.5% paraformaldehyde. Cells with and without Nocodazole were then re-plated on fibronectin in presence and absence of Nocodazole and fixed with 3.5% PFA after 5 mins. All the time points were immunostained with GM130, fixed, mounted and imaged using a confocal microscope.

### Arf1 activity assay

WTMEFs serum starved with 0.2% FBS containing DMEM for 12 hours were detached using Trypsin-EDTA, held in suspension for 120 mins (120’ SUSP), re-plated on fibronectin (10 µg/ml) for 15 mins (15’FN) and for 4 hours to be stable adherent (SA). Cells were lysed and processed for Arf1 activity assay as described earlier(Balasubramanian et al., 2007b). 30 µL of WCL and all of the GGA3 pulldown sample eluted with 20 µl of 2X lamelli buffer were resolved by 12.5% SDS-PAGE gel and transferred to PVDF membrane (Millipore). Blots were blocked with 5% milk in Tris-buffered saline containing 0.1% Tween-20 (TBST) for 1 hour at RT and incubated at 4°C overnight with the anti-Arf1 antibody (Clone 1D9, Abcam) diluted 1:500 in 2.5% milk in TBST. Blots were washed and incubated with anti-mouse HRP diluted 1:10000 in 2.5% milk in TBST at RT for an hour and developed with PICO chemiluminescence detection system (Thermo Scientific). LAS4000 (Fujifilm-GE) was used to image the blots and densitometric band analysis was done using Image J software (NIH).

To determine Percentage Arf1 activity, following calculation was used:

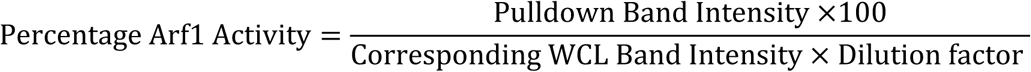

Dilution factor was calculated as the ratio of the amount of total cell lysate used for the pulldown (400 μl) and the amount of this lysate loaded as representative of whole cell lysate (WCL) on SDS PAGE, (22.5 μl WCL + 7.5 μl 4× Lamellis). The dilution factor was hence 400÷22.5=17.77. This ratio was kept constant in all the experiments in this study. Percentage active Arf1 levels thus calculated were compared between stable adherent, suspended and re-adherent cells.

Arf1 activity assay for inhibitor studies

For inhibitors studies (BFA, Ciliobrevin, and Nocodazole) percentage active Arf1 values for suspended or re-adherent cells were normalized to the control untreated cells and represented. Similarly, total Arf1 levels (Arf1/Actin in WCL samples) in inhibitor-treated cells were normalized to untreated controls and represented in every experiment.

### Immunofluorescence staining of cells for GM130

Serum-starved cells held in suspension as described earlier or re-adherent were fixed with 3.5% paraformaldehyde (PFA), permeabilized with PBS containing 5% BSA and 0.05% Triton-X100 for 10 mins at RT. Cells were then blocked with 5% BSA in PBS for 30 mins at RT and incubated with anti GM130 antibody diluted 1:100 in PBS with 5% BSA for an hour at RT. Cells were washed with PBS and incubated with 1:1000 diluted anti-mouse Alexa 488/ Alexa 568 antibody (as required) at RT for one hour. Suspended cells were similarly labeled in Eppendorf tubes and mixed regularly by tapping to prevent cells from settling. Cells were washed with 1X PBS and eventually reconstituted, mounted using flouramount-G and imaged using a confocal microscope.

### Confocal Microscopy

Cells were imaged using the Zeiss 710 or 780 laser scanning confocal microscopes using a 63X oil objective (NA 1.4). Acquisition settings were kept constant at: laser power = 2%, Pinhole = 1 AU, gain = 700 to 800 and images acquired at 1024 x 1024 resolution. Z-stacks were acquired at 0.2 µm interval thickness, deconvoluted and rendered as discussed below.

### Deconvolution of Z stacks and object analysis

All the images were processed and analyzed using the Huygens Professional version 16.10 (Scientific Volume Imaging, The Netherlands, http://svi.nl). De-convolution of confocal z-stack was optimised using the following settings. Average background value = 1, number of iterations = 30, signal to noise ratio (SNR) = 20, and quality change threshold = 0.0001. These settings were kept constant for all de-convoluted images discussed in this study. Theoretical point spread function (PSF) values were estimated for each z-stack and provide the minimal voxel size that the confocal microscope could resolve. This PSF value was then used in the software as garbage volume for surface rendering and object analysis. De-convoluted images were rendered either as a 3D maximum intensity projection (using the MIP Renderer plug-in) or surface rendered (using the Surface render plug-in with a 15% primary threshold). The top view of a MIP or surface rendered cell (zoomed 1.5X or 2.5X) was used to represent the Golgi phenotype. When needed a cross-sectional view along the Z-Axis of the MIP image was used to observe the localization of the Golgi in a re-adherent cell.

The number of discontinuous Golgi objects in a 3D deconvoluted image was determined using the advanced object analysis plugin in the Huygens Professional software. The garbage volume was set as calculated earlier, and a 15% threshold used to determine the number of discontinuous Golgi objects present in a cell. This was done for all cells in each treatment and the average number of Golgi objects determined. These were then compared between treatments to comment on the extent of the Golgi disorganization. Total Golgi volume was measured by addition of the Golgi volume of each object in a cell at a given time or treatment.

### Determining the Golgi distribution profile in a cell population

Cells were imaged using a confocal microscope and the structure of the Golgi observed and classified as organized or disorganized. Representative cross-sectional confocal images of organized and disorganized Golgi from each population or treatment were acquired, deconvoluted, surface rendered. A minimum of 50 (for bead experiments) and maximum of 200 randomly selected cells were observed in each population or treatment and their Golgi structure classified as above. These numbers we then used to calculate percentage distribution of organized vs disorganized Golgi in each population for a given time point or treatment.

### Co-localization analysis

Co-localization analysis for a cis Golgi marker (GM130) and ER marker (KDEL-RFP) was done using the Colocalization Analyzer Plug-in in the SVI Huygens Professional software (version 16.10). De-convoluted cross-section images of the cell expressing both markers were opened with this plugin and Pearson coefficients were calculated for each cell. Average values were then compared between suspended and re-adherent cells. A line plot of the intensities for each marker was made using the Huygens twin slicer plugin. Fluorescence intensities thus obtained for both markers were plotted using GraphPad Prism and the overlap in their intensities compared using the plot.

### Detection of Dynein bound with Active Arf1

Active Arf1 was pulled down using the GST-GGA3 construct as described earlier for the activity assay. Pull down and whole cell lysate (WCL) samples were probed for mouse anti-Arf1 antibody (clone 1D9, Abcam, Cat. No. # ab2806) at a dilution of 1:500 and mouse anti Dynein antibody (clone 74.1, Millipore, Cat. No. # MAB1618) at a dilution of 1:2000. Arf1 and dynein levels in the pulldown were normalized to respective levels in the WCL, values thus obtained normalized to stable adherent cells and compared suspended and re-adherent cells.

Serum-starved 10×10^5^ live WT MEF cells either treated with BFA (10 µg/mL in MeOH) or MeOH as the control for 2 hours. Cells were lysed and immediately processed for pulldown of Active Arf1 using GST GGA3 beads as described earlier. Active Arf1 levels and dynein bound in pull-down fractions were detected by western blot and compared.

### Cell membrane lectin labeling and quantitation by flow cytometry

For studies on lectin binding, WTMEFs serum starved for 12 hours were detached using Accutase (Sigma), processed and then held in suspension for 120 mins. 10×10^5^ live cells following detachment (5’ SUSP) and after 120min in suspension, were incubated with Alexa-488 conjugated ConA (0.025 µg/µL), PNA (0.025 µg/µL), HPA (0.025 µg/µL), WGA (0.0005 µg/µL) and FITC-UEA (0.1 µg/µL) for 15 mins on ice in the dark in 200 µl PBS. Cells were gently mixed during incubation to avoid clumping. They were eventually washed with cold PBS, fixed with 3.5% PFA for 15 mins at RT and re-suspended in 200 µL PBS. Cells were analyzed using the BD LSRFortessa SORP cell analyzer. Unlabelled detached (5 min) and 120 mins suspended cells were analyzed to set forward and side-scatter profiles and this was then used to set a gate to classify autofluorescence and then lectin fluorescence was measured. 10000 events were recorded for each treatment and time point to obtain levels of bound lectins on the cell surface. This data was analyzed using the Flowing software 2.5.1 and median fluorescence intensity was calculated for each cell population. Median fluorescence intensities of 120 mins suspended cells were normalized to their respective 5 min suspension time points and compared.

WTMEFs transfected with empty CherryN1, WT-Arf1-CherryN1 and Q71L-Arf1-CherryN1 using LTX-PLUS transfection reagent (Invitrogen) were serum starved for 12 hours, detached (5’ Susp) using Accutase and held in suspension for 120 mins. 10×10^5^ live cells (for each construct) were collected at both time points. Cells were surface labeled with Alexa-488 conjugated ConA (0.025 µg/µL), and FITC conjugated UEA (0.1 µg/µL) as described above. 7000 cells in the population gated for cherry fluorescence (Arf expression) were analyzed for lectin binding (Alexa 488/FITC). Flowing software 2.5.1 was used to determine median fluorescence intensity for bound lectin in cells suspended for 120 mins normalized to its respective 5 min suspension time point. Basal lectin levels (at 5 min suspension) were also compared across cells expressing Arf1 cherry constructs, as were the levels of cherry fluorescence levels for WT-Arf1-Cherry and Q71L-Arf1-Cherry constructs.

### Statistical analysis

All the analysis was done using Prism Graphpad analysis software. Statistical analysis of data for the number of Golgi objects, Arf1 activity, Pearson coefficient, and bound lectin levels were all done using the two-tailed unpaired Mann-Whitney t-test. When data were normalized to a control and compared, two-tailed single sample t-test was used. Statistical analysis for changes in distribution profile of Golgi phenotype was done using the two-tailed, Chi-square t-test.

